# Multi-site Ultrashort Echo Time 3D Phosphorous MRSI repeatability using novel Rosette Trajectory (PETALUTE)

**DOI:** 10.1101/2024.02.07.579294

**Authors:** Seyma Alcicek, Alexander Richard Craig-Craven, Xin Shen, Mark Chiew, Ali Ozen, Stephen Sawiak, Ulrich Pilatus, Uzay E Emir

## Abstract

**Purpose:** This study aims 1) to implement an operator-independent acquisition, reconstruction, and processing pipeline using a novel rosette k-space pattern for UTE ^31^P 3D MRSI and 2) to evaluate the clinical applicability and replicability at different experimental setups.

**Methods:** A multicenter repeatability study was conducted for the novel UTE ^31^P 3D Rosette MRSI at three institutions with different experimental setups. Non-localized ^31^P MRSI data of 5 healthy subjects at each site were acquired with an acquisition delay of 65 μs and a final resolution of 10 × 10 × 10 mm^3^ in 9 min. Spectra were quantified using the LCModel package. The potential acceleration was achieved using compressed sensing on retrospectively undersampled data. Reproducibility at each site was evaluated using the inter-subject coefficient of variance.

**Results:** This novel acquisition and advanced processing techniques yielded high-quality spectra and enabled the detection of the critical brain metabolites at three different sites with different hardware specifications. In vivo, feasibility with an acceleration factor of 4 in 6.75 min resulted in a mean Cramér-Rao lower bounds below 20% for PCr, ATPs, PME, and the mean CoV of ATP/PCr resulted in below %20.

**Conclusion:** We demonstrated that UTE ^31^P 3D Rosette MRSI acquisition, combined with compressed sensing and LCModel analysis, allows patient-friendly, operator-independent, high-resolution ^3**1**^**P** MRSI to be acquired at clinical setups.

## Introduction

Phosphorous (^31^P) Magnetic Resonance Spectroscopy (MRS) has shown much promise as a non-invasive molecular imaging tool since the earliest high-resolution Nuclear Magnetic Resonance (NMR) spectra were obtained from phosphate-containing metabolites in the early 70s ^1,2^. The first non-invasive molecular imaging followed these efforts via ^31^P MRS^3^ and its utilization in disease^4^, wherein the technique provides a wide variety of molecular information. This includes adenosine triphosphate (ATP) and phosphocreatine (PCr), markers for deficits in energy metabolism, phosphomonoesters (PME, phosphorylethanolamine (PE) + Phosphorylcholine (PC)) and phosphodiesters (PDE, GPE+GPC), markers for cell synthesis and myelination, and nicotinamide adenine dinucleotide (NADH), a marker for cellular redox status ^5,6^. Thus, ^31^P MRS can contribute to diagnosing and monitoring disease before structural/extracellular changes become apparent and can also allow measuring modulations in metabolites during physiological interventions. Alterations in these metabolites indicate impairment in energy storage and membrane synthesis or breakdown^7–9^. Recent studies using ^31^P MRS have reported alterations in energy metabolism metabolites in AD and MCI compared to healthy controls, interpreted as dysregulation of energy metabolism^10,11^. In addition, changes in ^31^P MRS-derived PME and PDE composition have also been reported in individuals at high risk for AD, reflecting the myelin and proliferation-related alterations ^12^. Furthermore, similar to the ^1^H MRS cases^13^, it has been demonstrated that phospholipid metabolites of ^31^P MRS might identify the status of mutation of the isocitrate dehydrogenase in patients with glioma^14^. As importantly, in addition to the energy and phospholipid metabolism, ^31^P MRSI shed light on the intracellular pH. Recently, it has been demonstrated that intracellular pH alteration might be linked to tumor proliferation^15^.

^31^P MRS imaging (MRSI) offers metabolic profiles over larger regions, allowing shorter acquisition delays in contrast to fully localized acquisitions. Alternatively, to avoid the lengthy slice-selective gradients for localized acquisitions, outer volume suppression (OVS) methods such as image-selected in vivo spectroscopy (ISIS) have been the method of choice for ^31^P MRS^16^. However, lower sensitivity, limited spectral-spatial resolution, and prolonged acquisition durations hamper the broader application of ^31^P MRSI in the clinical environment^5^. Also, commonly used acquisition delays (TE>300 μs) with conventional methods result in phase and baseline roll and are subject to operator errors in metabolite quantification^5^. Time domain fitting algorithms have been used to overcome this problem^17^.

As increasingly advanced MRSI acquisition technologies have become available^18^, their improved resolution at comparable sensitivity can benefit ^31^P MRSI acquisition and quantification and facilitate robust clinical application of MRSI techniques. For instance, accelerated kspace trajectories ^19–21^, improved reconstruction approaches ^20,22^, and increased SNR with the UHF^23^ have mitigated long acquisition durations and improved SNR and resolution. In addition, several offline data analysis tools have demonstrated the feasibility of operator-independent metabolite quantification of data with phase distortions and baseline roll ^24,25^.

This work intends to develop a novel acquisition technique at 3T that addresses the challenges of speed, spatial resolution, first-order phase, and baseline roll of ^31^P MRSI. Thus, this study has implemented a novel rosette k-space pattern for UTE ^31^P 3D MRSI^26^ and assessed it by measuring ^31^P metabolites. Further acceleration with compressed sensing (CS) is demonstrated with retrospectively undersampled rosette trajectory. The performance of the proposed operator-independent acquisition, reconstruction, and processing pipeline is assessed with LCModel analysis. Finally, the clinical applicability and replicability of UTE ^31^P 3D Rosette MRSI readouts at different experimental setups are evaluated using inter-subject reproducibility at three different institutions.

## Methods

This multicenter reproducibility study was conducted in accordance with the regulations of the local institutional human ethics committees. Five healthy volunteers were scanned. The mean ± standard deviation (SD) age of the volunteers was 24 ± 5 years (2 females), 35 ± 3 (1 female), and 30 ± 2 (2 females), respectively.

### Institutions and hardwares

Volunteers underwent brain scans with a whole-body 3T MRI system (Siemens, Erlangen, Germany). The contributing institutes and corresponding scanner configurations were:

**Site 1** Purdue University, USA: Magnetom Prisma with a maximum gradient amplitude of 80□mT/m and maximum slew rate of 200□mT/m/ms isotropically, Dual Tuned Surface Coil, RAPID Biomedical.
**Site 2** Haukeland University Hospital, Bergen, Norway: Biograph mMR with a maximum gradient amplitude of 40□mT/m and maximum slew rate of 180□mT/m/ms isotropically, Dual Tuned Quadrature Head Coil, RAPID Biomedical.
**Site 3** University Hospital Frankfurt, Goethe University, Germany: Magnetom Prisma with a maximum gradient amplitude of 80□mT/m and maximum slew rate of 200□mT/m/ms isotropically, Dual Tuned Quadrature Head Coil, RAPID Biomedical.

At all sites, the transmit voltage of the system was optimized to achieve a desired excitation flip angle by maximizing the transverse magnetization M_xy_ for a 90° flip angle on the X-nuclei Transmit Voltage card of the vendor-provided software. The vendor provided ^1^H B_0_ shimming procedure was performed in a volume of 240^3^ mm^3^ for Site 1, 70×100×480 mm^3^ for Site 2 and 60^3^ mm^3^ for Site 3.

### UTE ^31^P 3D Rosette MRSI acquisition

A vendor-provided multi-plane isocenter localizer was performed for anatomical reference, followed by a UTE ^31^P 3D Rosette MRSI acquisition. The parameters used in the UTE ^31^P MRSI sequence with rosette k-space sampling were: Field of view = isotropic 480×□mm^3^ for Site 1 and 3 (a maximum slew rate of 187.27 mT/m/ms) and 600^3^ mm^3^ for Site 2 (a maximum slew rate of 149.81 mT/m/ms), the readout dwell time = 5□μs, TR = 350□ms, readout duration = 275□ms, and RF pulse duration = 50□μs. This study used an Ernst flip angle, assuming that the T_1_ of PCr at 3T is 2.6s ^27^. The ADC with a bandwidth of 200 kHz was turned on 10 μs after RF, and readout gradients were turned on 30 μs after ADC, resulting in an acquisition delay of 65□μs. These extra 6 ADC (6 × 5□μs) points were discarded before reconstruction (Figure 1). Before the MRSI data acquisition, 100 dummy TRs were run to reach the steady state.

**Figure 1.**
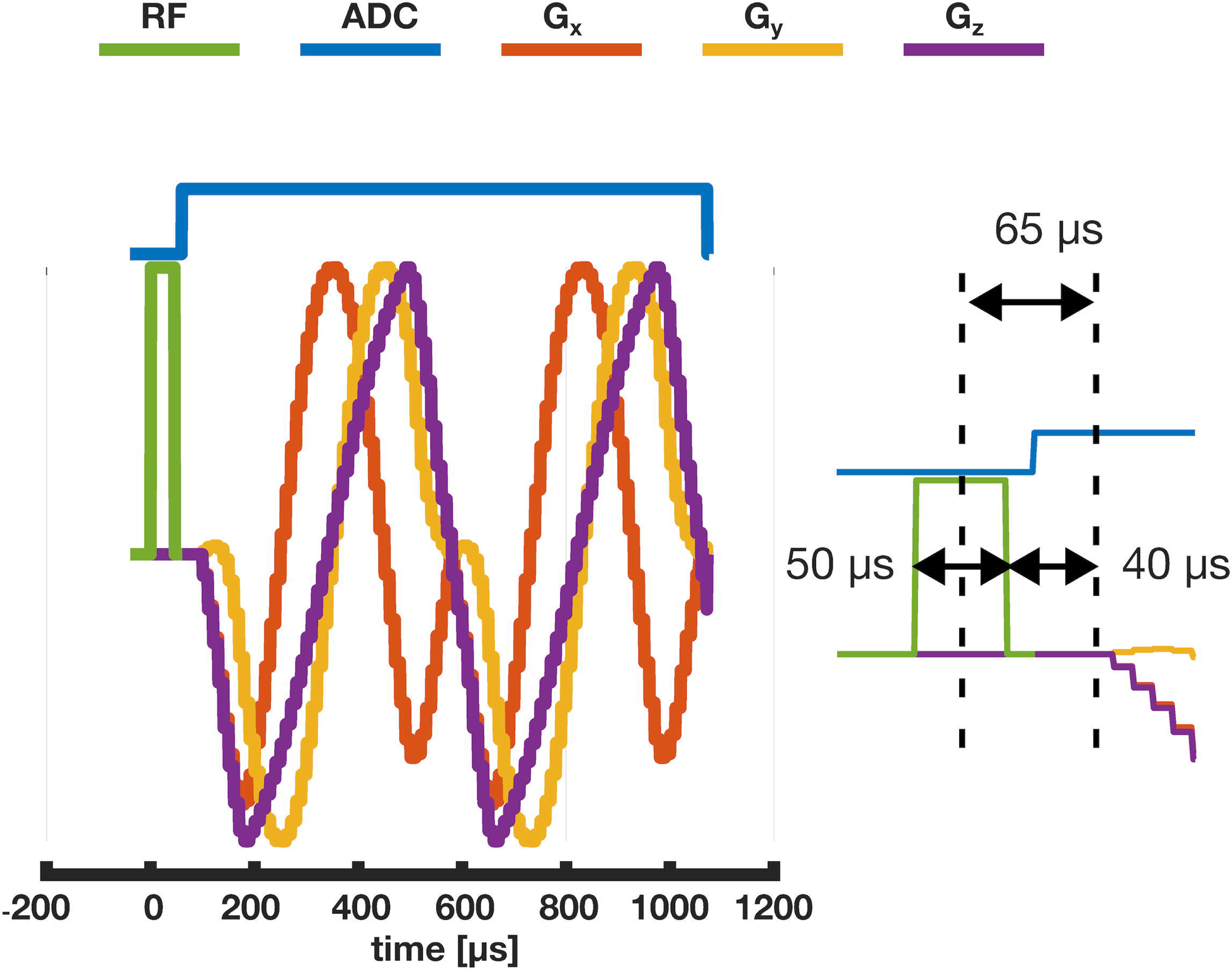
The detailed pulse diagram illustrates achieving an acquisition delay of 65 μs from the center of the RF pulse. ADC event starts ten us after the RF pulse. Readout gradients switch on after 6 ADC points (30 μs). The total delay for readout gradients from the end of the RF pulse is 40 μs.

The following equations were used for the 3D rosette MRSI k-space trajectory with a specific case where w_1_ and w_2_ are equal^28^.

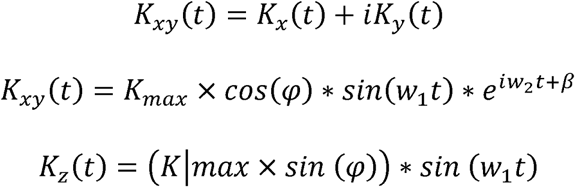

where *K_max_*is the maximum extent of k-space, w_1_ is the frequency of oscillation in the radial direction, w_2_ is the frequency of rotation in the angular direction, *φ* determines the location in the z-axis, and *β* determines the initial phase in the angular direction.

According to this, each rosette petal was designed with 96 points per rosette with w_1_=w_2_ of 6500 rad/s (*w*_1_ = *π*⁄(*dwelltime* × *numberofpointsperpetal*(*N_pp_*))). For the reconstruction, 96 points were downsampled to 48 by simply averaging the oversampled points, resulting in an effective bandwidth of 100 kHz. Since dual-echo images can be generated within a single acquisition with a manual separation at the middle of each data readout of the novel rosette acquisition, only the first half of the petal (*N_pp_*/2 = 24) was used for the reconstruction. According to Nyquist’s criteria, the required number of petals (*N_p_*) for a matrix size of 24 was calculated to be 1810 (4 × *π* × 12^2^). Due to the efficient sampling of the rosette k-space pattern, only 80% of the required k-space, 1444, can be considered full-k-space acquisition ^28^. The total acquisition time for UTE ^31^P 3D Rosette MRSI was around 9 min (Np × TR + dummy scans × TR= 505 + 35 s).

256 spectral points (number of petal repeat or spectral points, *N_sp_* = 256) were collected with an effective spectral bandwidth (SBW) of 2083 Hz for ^31^P MRS, corresponding to a spectral resolution of 8.1 Hz (Figure 2a,b). Crusher gradients in all three directions were applied at the end of each readout gradient and before each excitation pulse. The total acquisition time for UTE ^31^P 3D Rosette MRSI with three averages was 27 minutes.

**Figure 2.**
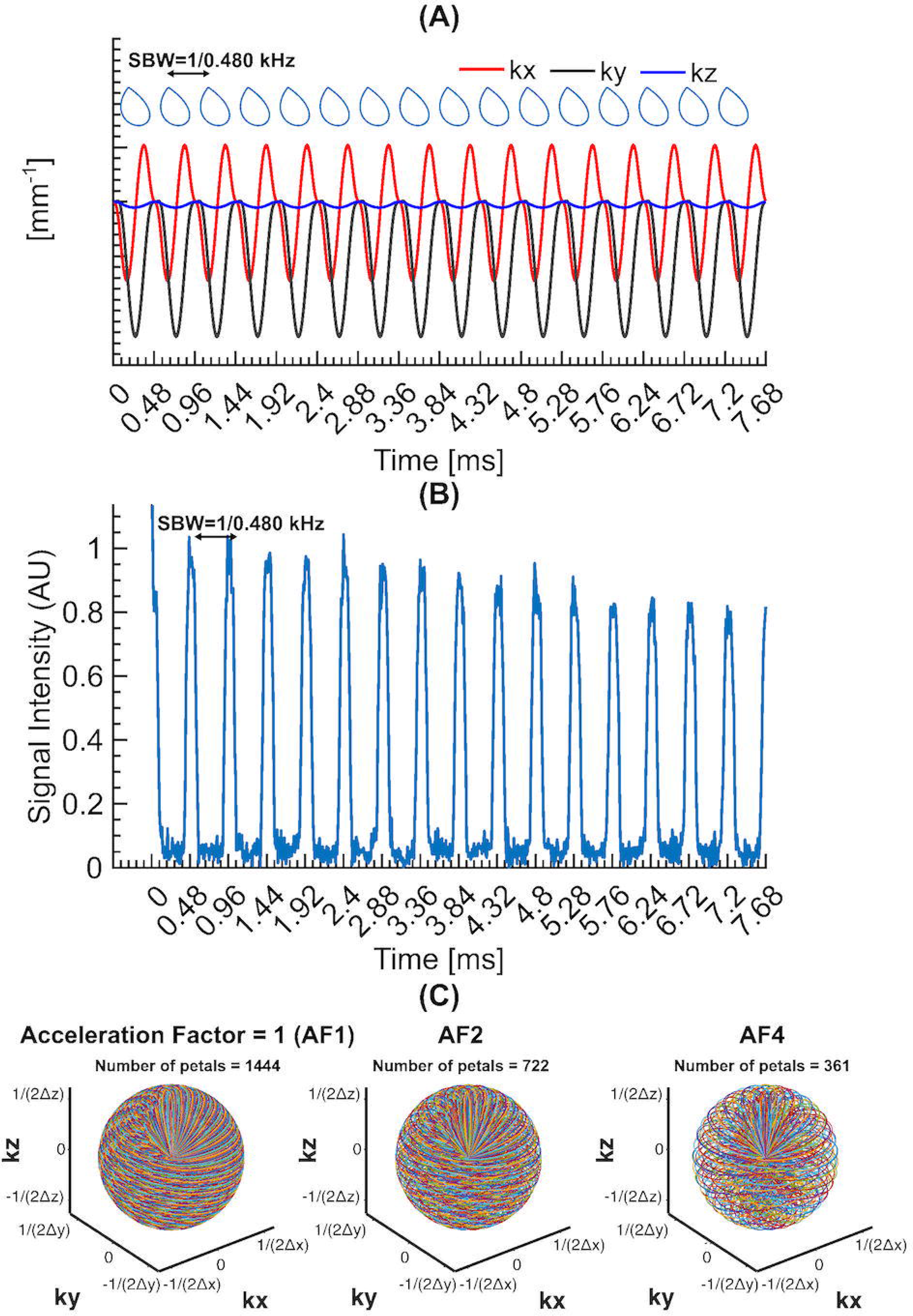
(A) The detailed k-space diagram illustrates the repetition of the same petal every 480 μs to achieve a spectral bandwidth of 2.083kHz. (B) A representative k-space data was acquired from a uniform phantom of inorganic phosphate. Only the first 16 repeats were illustrated for an average of 1444 petals. (C) An illustration of the 3D rosette k-space design with different acceleration factors (AF1, AF2, and AF4).

### Reconstruction and Pre-processing

Image reconstruction and post-processing steps were performed in MATLAB (MathWorks, USA). The nonuniform fast Fourier transform was used to calculate the forward encoding transform of the acquired k-space data ^29^. A CS approach was used for image reconstruction, using total generalized variation as the sparsifying penalty^30,31^.

Specifically, NUFFT calculated the Fourier transform of the acquired 3D *k*-space MRSI data (*k_xy_* × *k_z_* = 48 × 48, and N_pp_ × N_sp_ = 24 × 256) on the trajectory for all time frames to generate 4D MRSI data with a final matrix size of 48×48×48×256. Retrospective undersampling of fully sampled datasets with compressed sensing was also analyzed for acceleration factors (AFs) of 2 and 4 for prospective acceleration (Figure 2c). The resulting FIDs for all AFs were zero-filled to 1024 points and filtered with a Gaussian filter of 100 ms, a 1 Hz line broadening function, and zero-order phase-corrected.

### Metabolite Level and Spectral Quality Metrics

The LCModel package was used to quantify the metabolite spectrum for each MRSI voxel ^32^. The model spectra of PCr, α-ATP, β-ATP, γ-ATP, inorganic Phosphate (P_i_), PE, PC, GPE, GPC, nicotinamide adenine dinucleotide (NADP, reduced form, NADH and oxidized form, NAD+), membrane phospholipids (MP) and 2,3-diphosphoglycerate (DPG) were simulated using in-house MATLAB codes with published ^31^P chemical shifts and J-coupling constants ^24^. Simulations with an ideal excitation pulse were performed using the same sequence timings (SBW and acquisition delay) as those on the 3□T system in use (Figure 1). Since the quantifications were not corrected for T_1_ and B_1_ due to the scan time constraints, we therefore chose to express results as metabolite ratios, PCr/ (α-ATP+β-ATP+γ-ATP) (PCr/ATP), and phosphomonoesters (PME)/phosphodiesters (PDE). In addition, to compare the performance of 3D MRSI at different sites, the SNR (SNR_LCModel_) and linewidth (LW_LCModel_) estimations of LCModel were used as spectral quality metrics. SNR_LCModel_ was calculated using the peak height of the PCr singlet peak and the root-mean-square of its residual, and LW_LCModel_ was calculated from the width of the three peaks defined in the LCModel control file (PCr, P_i_, NADP). The control file for the LCModel fitting is provided in the supporting information (Figure 1S).

### Regional Distributions of ^31^P Metabolites and Statistical Analysis

The resulting MRSI resolutions were isotropic 10 mm^3^ for Sites 1 and 3 and 12.5 mm^3^ for Site 2. Each subject’s MRSI slice and metabolite maps were determined in the subject’s image space. The Montreal Neurological Institute-152 (MNI) template was downsampled to 10 mm^3^ to match the MRSI resolution for coregistration purposes. Each subject’s MRSI volumes of Site 2 and 3 were coregistered to the MNI template using FMRIB’s Linear Image Registration Tool (FLIRT) implemented in the FMRIB’s Software Library (FSL) ^33^. Since the surface coil used in Site 1 resulted in limited brain coverage (occipital cortex), each subject’s MRSI volume from Site 1 alignment to MNI space was manually conducted using FSLeyes nudge ^34^. Afterward, mean metabolite levels in white matter and gray ROIs were calculated using fslstats^33^. Due to the different performances of the surface and volume coils used in this study, two sets of gray matter and white ROIs were identified for mean metabolite levels (Figure S2). The workflow of post-processing steps was summarized in Figure S3.

For each site, the inter-subject variance was measured separately through the Coefficient of Variance (CoV). This analysis was performed on each voxel where PCr had Cramér-Rao lower bound (CRLB) (goodness of fit) values smaller than 20%. Metabolite ratios and spectral quality metrics for all voxels within an ROI were averaged together to calculate the ROI-specific values (Figure S2). The interactive AFNI view, showing slices and spectra graphs, was used to visualize the final results spectra^35^. Mean and standard deviation were determined from ROI-averaged metabolic ratios for every subject. Site-specific inter-subject values were then averaged to calculate the inter-subject CoVs. Due to the different hardware setups, our acquisition protocol did not account for any statistical comparison between sites.

As for reproducible research, we provide some MATLAB scripts and k-space data to reproduce some of the results described in this article. The software and data can be downloaded from https://purr.purdue.edu/projects/ismrm31pmrsi.

## Results

Figure 3 shows the UTE ^31^P 3D Rosette MRSI spectra on a 5 × 3 grid, with LCModel fits together with their first-time point of FID images. Due to hardware specifications, Site 1 resulted in coverage limited to the occipital cortex, while full brain coverage was achieved for Sites 2 and 3. The first-time point FID images generated images with structural information for sites 2 and 3. Even at a final resolution of 10 mm^3^ at 3T, spectra from 3 sites with different hardware setups are of sufficiently high quality for reliable LCModel analysis with full-kspace and retrospective AFs. The UTE ^31^P 3D Rosette MRSI with an acquisition delay of 65 μs and processing pipeline with LCModel parameters results in a linear baseline and zero-order phase of the maximum of 25 degrees (Figure 4 and Figure S4). As illustrated, similar baseline and phase distortions are also achieved with increased AFs while noise level rises due to undersampling.

**Figure 3.**
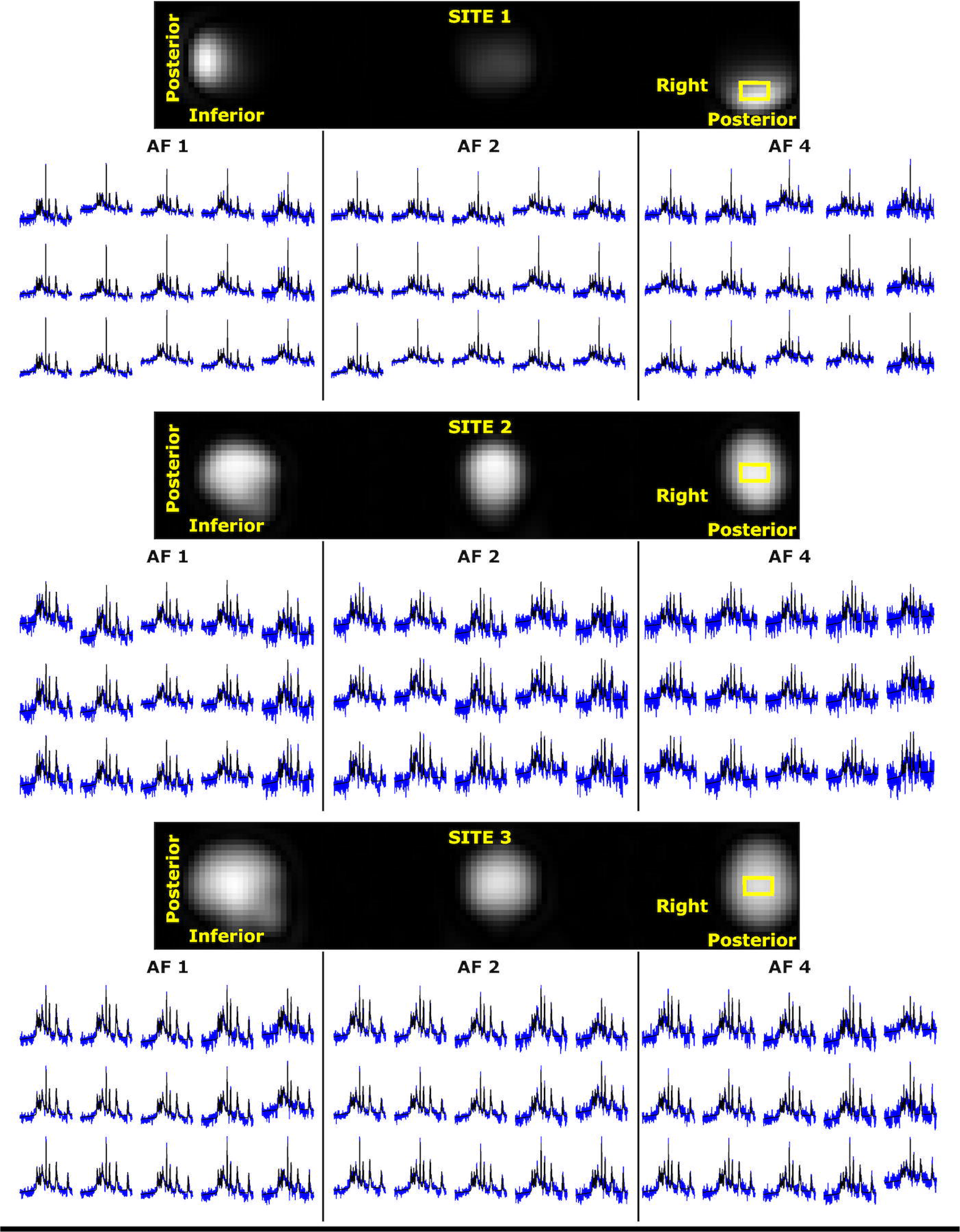
Extracted in vivo spectra (blue) from 15 voxels of a 5×3 grid with LCModel fit (black) for AF1, AF2, and AF4. The location of the 5×3 was illustrated as a yellow box on the first-time point of FID images.

**Figure 4.**
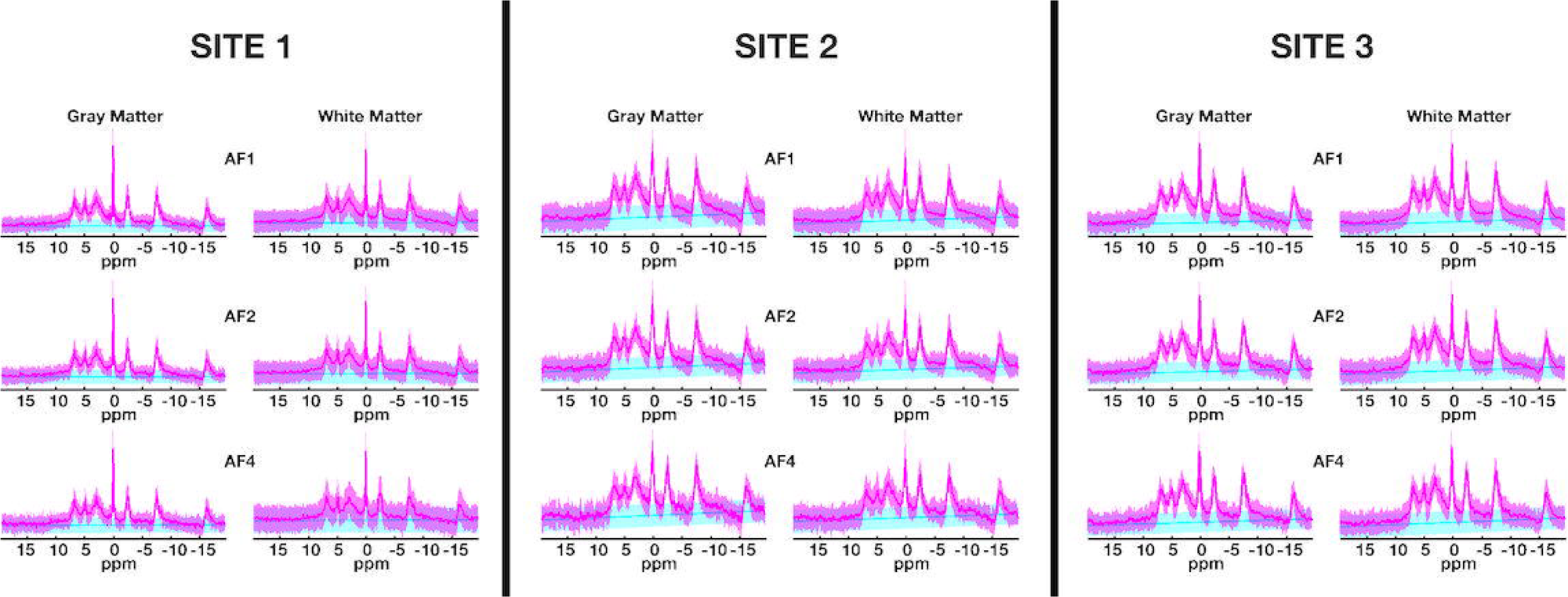
Mean (solid line) and ± standard deviation (shade) of ^31^P MRS mean spectra (magenta) and LCModel baseline estimations (blue) in each ROI from all subjects for different AFs. Each column separated by black lines indicates different sites.

The mean Cramer-Rao Lower Bound (CRLB) values of four reported metabolites (PCr, ATP, PME, and PDE) are lower than 25% in each ROI across subjects (Figure S5). Notably, the standard deviations of PDEs’ CRLBs are higher than the others, indicating PDE might not be reliably detected.

Figures 5 and 6 show high-resolution (10 mm^3^) metabolite ratios, spectral quality metrics, and corresponding inter-subject CoVs maps for all AFs for sites 1 and 3, respectively. The mean metabolite ratios for each ROI across subjects for each site and AFs are reported in Figure 7. Overall, mean metabolite ratios of PCr/ATP and PME/PDE resulted in similar values across sites and ROIs. This does not change with increased undersampling (AFs). For instance, mean PCr/ATP in gray and white matter across subjects and sites for full-kspace acquisition are 0.38±0.02 and 0.39±0.02, respectively, whereas mean PME/PDE in gray and white matter are 1.49±0.18 and 1.50±0.25, respectively. Similar values are also observed for AF of 4, where mean PCr/ATP in gray and white matter were 0.38±0.02 and 0.39±0.03, respectively, whereas mean PME/PDE in gray and white matter were 1.49±0.18 and 1.50±0.25, respectively.

**Figure 5.**
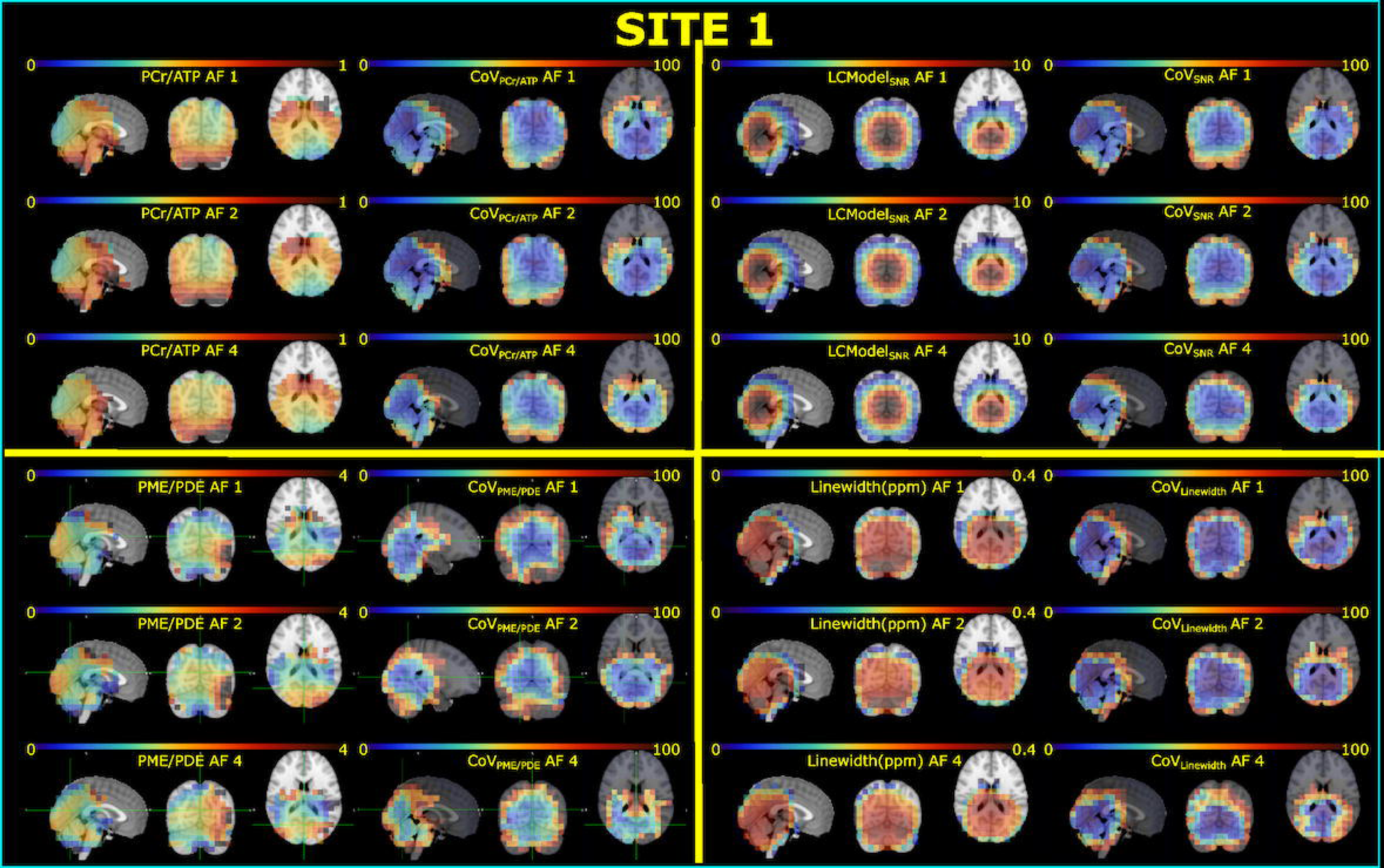
Metabolite ratios (PCr/ATP and PME/PDE), spectral quality metrics (SNR_LCModel_ and Linewidth (LW)_LCModel_), and corresponding inter-subject CoVs maps for all AFs for sites 1. Maps are overlaid on the Montreal Neurological Institute-152 (MNI) template. Voxels resulting in a CoV higher than 100 % were masked for each map.

**Figure 6.**
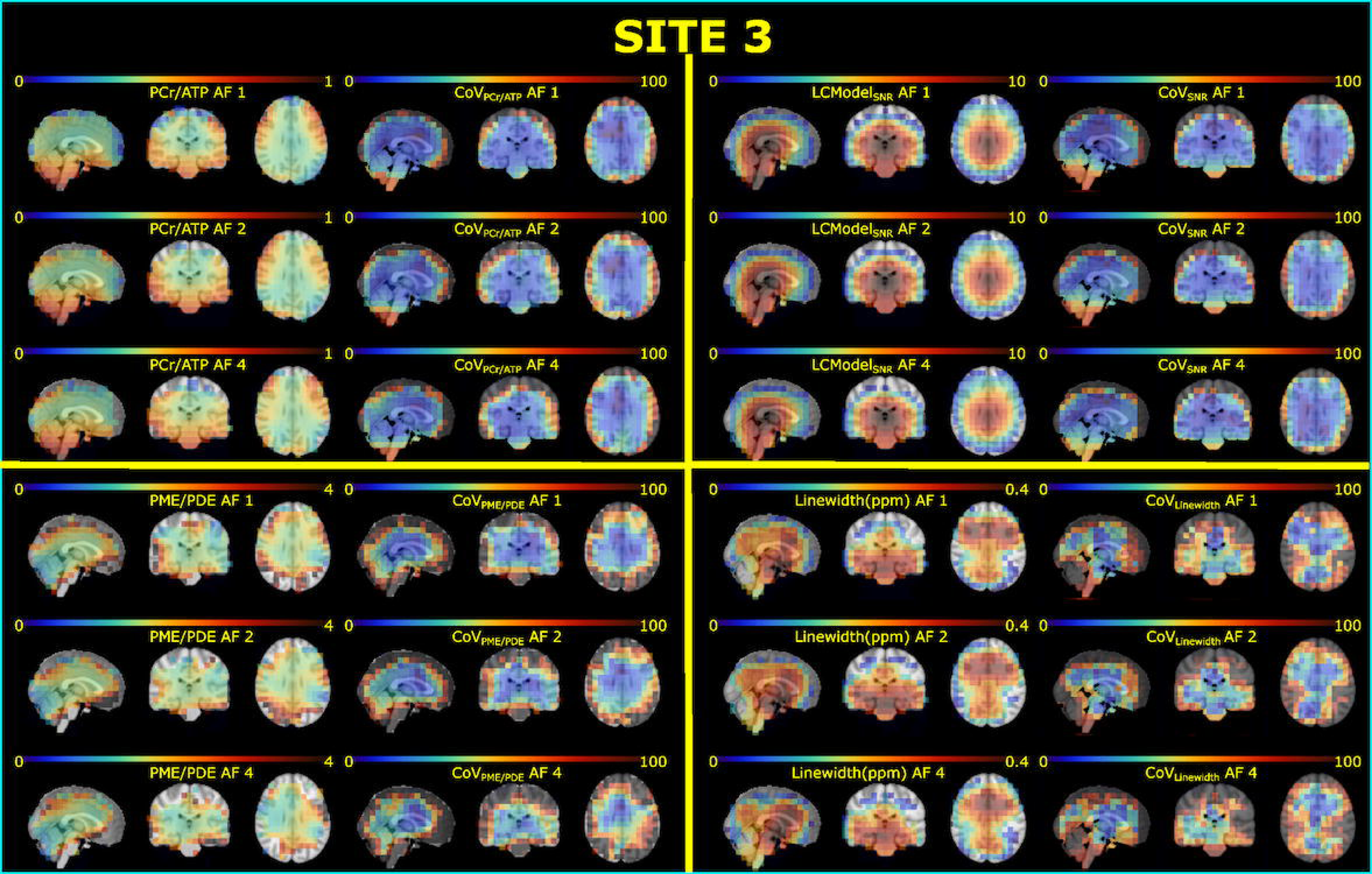
Metabolite ratios (PCr/ATP and PME/PDE), spectral quality metrics (SNR_LCModel_ and Linewidth (LW)_LCModel_), and corresponding inter-subject CoVs maps for all AFs for sites 3. Maps are overlaid on the Montreal Neurological Institute-152 (MNI) template. Voxels resulting in a CoV higher than 100 % were masked for each map.

**Figure 7.**
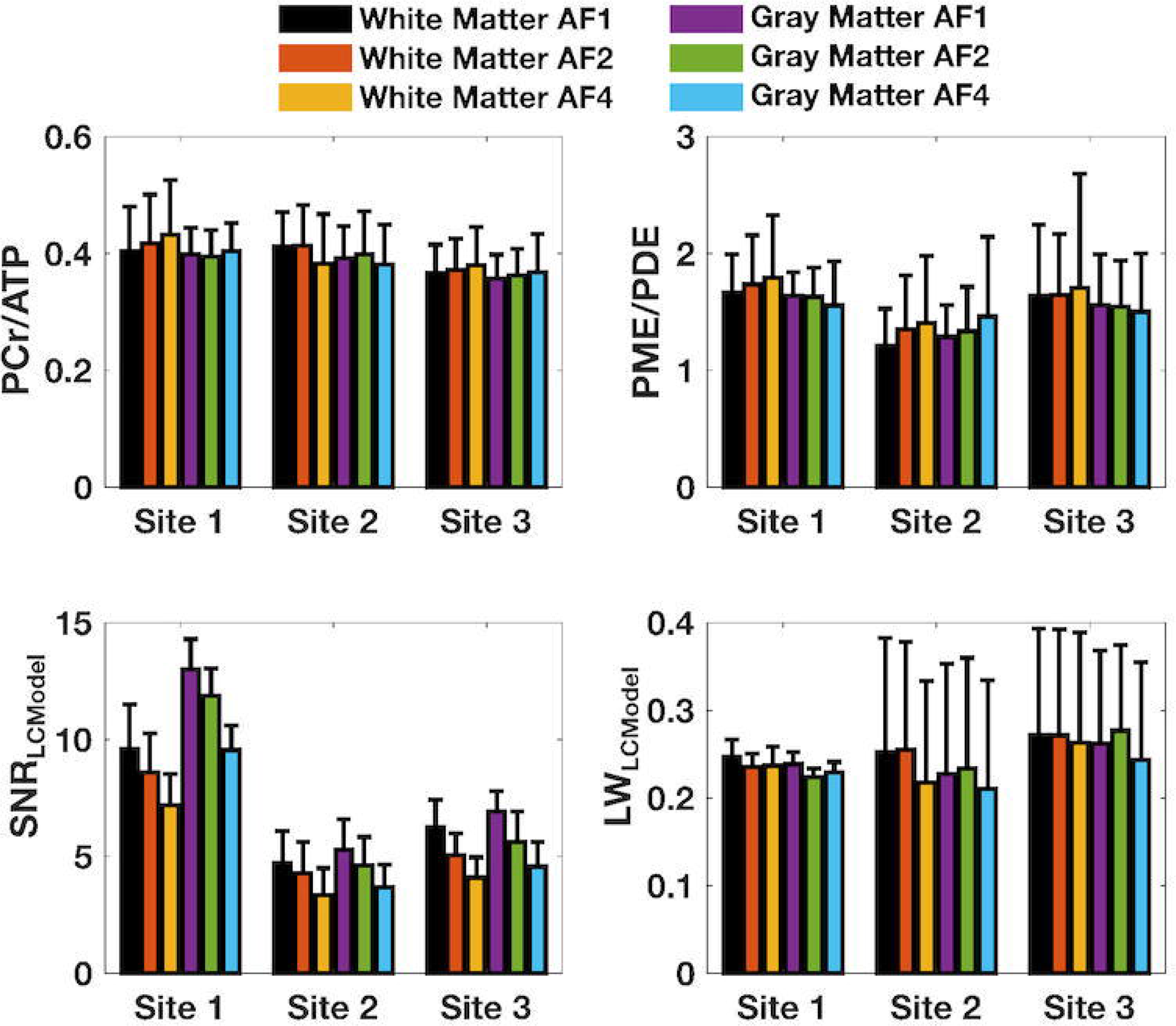
The mean metabolite ratios (PCr/ATP and PME/PDE) and spectral quality metrics (SNR_LCModel_ and Linewidth(LW)_LCModel_) for each ROI across subjects for each site and AFs.

As for spectral quality metrics, there are significant performance differences between sites due to different setups. The mean SNR_LCModel_ analysis reveals that the surface coil (Site 1) has higher values compared to volume coil findings for all AFs. As expected, the SNR_LCModel_ decreased with increased undersampling for all sites. LW_LCModel_ findings reveal the effect of effective shim volume due to the different receive sensitivity of the coils used across sites. However, the mean LW_LCModel_ values are similar across sites where sites utilizing volume coil (Site 2 and 3) result in higher standard deviations.

The mean inter-subject CoV for metabolite ratios for each ROI and AFs are reported in Figure 8. As in metabolite ratios, overall, CoV of metabolite ratios of PCr/ATP and PME/PDE resulted in similar values across sites and ROIs. CoVs for metabolite ratio increased with AFs. For instance, the mean inter-subject CoV of PCr ATPs in gray and white matter of full-k-space acquisition for all sites are 11.22±1.67 and 13.45±1.22, respectively, whereas mean PME/PDE in gray and white matter are 18.75±8.02 and 22.02±3.58, respectively. Degraded CoVs are observed for AF of 4, where mean PCr ATPs in gray and white matter were 15.89±3.32 and 18.90±3.11, respectively, whereas mean PME/PDE in gray and white matter were 29.96±5.21 and 35.17±8.32, respectively.

**Figure 8.**
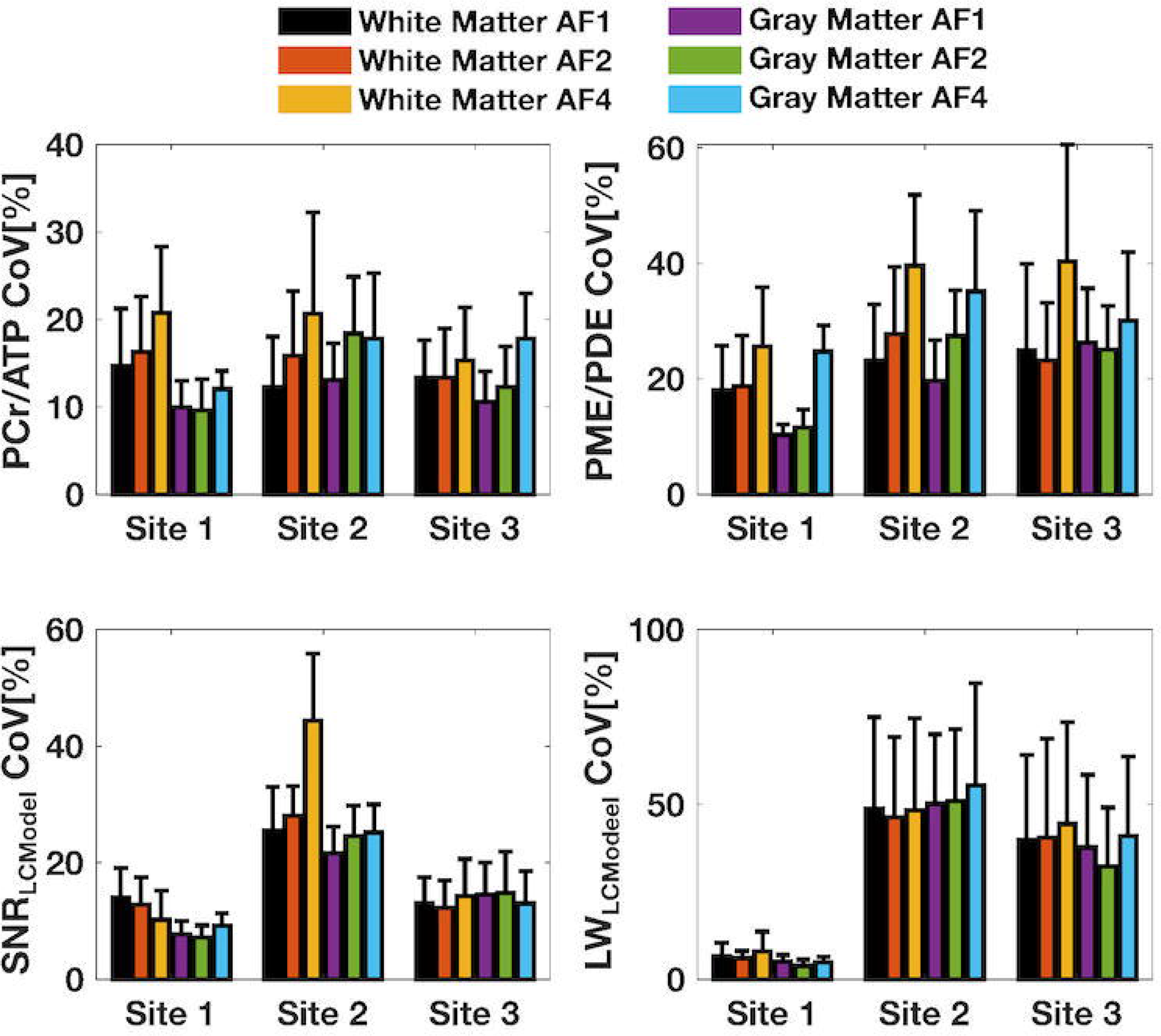
The mean inter-subject CoVs of metabolite ratios(PCr/ATP and PME/PDE) and spectral quality metrics(SNR_LCModel_ and Linewidth(LW)_LCModel_) for each ROI across subjects for each site and AFs.

As for spectral quality metrics, as in the mean SNR_LCModel_ and LW_LCModel_, the effect of different setups across sites echoed on CoVs of SNR_LCModel_ and LW_LCModel_. While CoVs remain lowest for Site 1, Sites 2 and 3 resulted in degraded CoVs for both SNR_LCModel_ and LW_LCModel_

## Discussion

This study developed and demonstrated a basic acquisition protocol with an automated reconstruction and LCModel analysis pipeline for ^31^P MRSI. In addition, we implemented an identical UTE ^31^P 3D Rosette MRSI sequence to assess reproducibility in healthy volunteers at three sites with different hardware specifications. This novel acquisition and advanced processing techniques yielded high-quality spectra and enabled the detection of the critical brain metabolites PCr, ATPs, PME, and PDE. In vivo feasibility with an AF of 4 in 6.75 min has been demonstrated with acceptable tradeoffs between scan time and spatial resolution, indicating the optimal spatial-temporal-spectral resolution using compressed sensing. High spectral quality is consistently achieved with negligible baseline and phase distortion due to the unprecedented short acquisition delay for MRSI acquisition on a clinical scanner. As such, this work initiates standardization efforts for the long-awaited ^31^P MRSI on clinical scanners. Together, these results demonstrate the suitability of the proposed UTE ^31^P 3D Rosette MRSI technique for clinical and experimental use.

### Novel Rosette And Compressed Sensing

A significant barrier against the wide use of ^31^P MRSI in clinical practice is the need for a prolonged acquisition to compensate for the lower abundance and lower sensitivity (correspondingly low SNR) of ^31^P. Accelerated acquisition trajectories have been developed ^18^ and are in place to reduce the acquisition time with advanced reconstruction methods^22^, including the compressed sensing developed at 3T^20^. However, these techniques require additional temporal and/or spatial interleaves, resulting in a longer acquisition duration to reach the required broad spectral dispersion of the spectrum due to the lower gyromagnetic ratio of ^31^P. Alternatively, a limited spectral range has been used to eliminate the temporal leaves, resulting in missing essential features of the ^31^P spectrum^19,21^. Furthermore, some of these efforts have been implemented at UHF to compensate for the SNR deficiency but still require the additional cost of temporal interleaves and/or limited spectral range to fulfill the broad spectral dispersion requirement of the ^31^P^19,21^. This study successfully demonstrated the accelerated ^31^P MRSI method using a modified rosette trajectory for brain applications^28^. The choice of rosette k-space trajectory resulted in significant improvements over other acquisition strategies. First, we successfully modified our novel rosette space trajectory to reach the required spectral bandwidth for ^31^P without requiring any spatial or temporal interleaves. In line with our earlier efforts, the proposed strategy has improved SNR per acquisition time compared to the radial k-space ^28^. Finally, as demonstrated earlier, the CS reconstruction performs better when the kspace is incoherently sampled more in the center ^28^. Therefore, we modified the rosette kspace trajectory to fulfill this by oversampling at the center k-space^28^. Using CS, we used this SNR improvement for higher-resolution reconstruction (10 mm^3^) and further acceleration, AF4. In contrast to earlier 3D ^31^P MRSI studies with acquisition duration ranging from 15 to 27 minutes with a resolution of ∼25 mm^3 5^, the proposed method with AF4 achieves 10 mm^3^ resolution in 6.75 minutes, making the clinical ^31^P MRSI patient-friendly.

### Unprecedented UTE

In addition to the low intrinsic signal-to-noise ratio (SNR) of ^31^P, the short T_2_ relaxation times of ^31^P metabolites, i.e., T_2 ATP_ ∼ 50 ms at 3T), acquisition strategies with minimal acquisition delay are preferable over fully localized acquisition strategies ^36^.

Depending on the choice of excitation pulse, minimal acquisition delay is typically around 300 us ^5,6^. In this study, the ADC was turned on 10 μs after the RF pulse, followed by readout gradients. Sampling during the gradient ramp-up period enabled the shortest possible acquisition delay of 65 μs. It provided immunity to short T_2_ relaxation times of ^31^P metabolites.

### Spectral Quality and LCModel fitting

Several approaches to quantifying in vivo MR spectra, including ^31^P, have been described in ^24,25^. In particular, automated analysis of many ^31^P spectra of low SNR from a typical 3D-MRSI acquisition is challenging. Due to the choice of RF pulse and phase-encoding gradients in conventional acquisition strategies, ^31^P spectra inherited a first-order phase and baseline distortions. Mitigation of these artifacts requires operator-dependent evaluations. This study utilized the LCModel approach for automated analysis of ^31^P MR spectra ^24^. Further success of the automated analysis originates from an unprecedented acquisition delay of 65 μs, resulting in minimal phase-roll and baseline distortions. Overall, using the LCModel with UTE ^31^P 3D Rosette MRSI resulted in automated evaluation, potentially minimizing potential methodological biases. For this, voxels were restricted to those with CRLB values of PCr below 20 % for the respective metabolite ratio and spectral metrics for all acceleration factors. Figure 4, S3 and S4 illustrate that regardless of the spectral quality degradation due to the retrospective undersampling the automated LCModel analysis resulted in similar baseline and zero-order phases without operator manipulation. Thus, an automated voxel selection protocol was successfully implemented for all sites with different AFs.

### Metabolite Ratios

To mitigate potential biases originating from different hardware setups and short TR (350 ms), we chose to report metabolite ratios. In general, metabolite ratios reported in this study agree between sites and those measured in previous studies. In particular, the PCr/ATP ratios measured in this study are in good agreement with those reporting PCR/total ATP (α-ATP+β-ATP+γ-ATP) and retrospective analysis of the ones that report either PCR/α-ATP or β-ATP ^10,20,24,37,38^. While similar mean PCr/ATP values are achieved with increased AFs, due to the noise amplification originating from undersampling, CoVs are increased. Similarly, PME/PDE values are in line between sites and with previous studies. In particular, this agreement holds firmly with studies that model the MP in their evaluations^10,24,37^. The Narrower chemical shift dispersion of ^31^P MRS at lower magnetic fields (3T:c) results in an overlapping between MP and PDE signals. If MP is not modeled in the quantification approach, this results in exacerbated PDE values, as reflected in the higher CRLB standard deviation of PDE (Figure XX), and reduced PME/PDE. Thus, the results of metabolite ratios must be cautiously interpreted. As in PCr/ATP, similar degradation was observed on CoVs for PME/PDE with AFs, while metabolite ratios hold similar values across sites. While the identical UTE ^31^P 3D Rosette MRSI acquisition method with automated reconstruction, spectral processing, and fitting were utilized, the CoV differences between the sites might suggest the different coil transmit profiles combined with the main magnetic field homogeneity achieved in the study.

### Limitations

While this study has demonstrated the feasibility and reproducibility of automated in vivo detection of ^31^P metabolites using novel accelerated acquisition and processing pipelines at three different hardware setups using 3T scanners from the same vendor, several factors affect the accuracy of the reported findings. First, achieving narrower linewidth is essential at 3T, where the chemical dispersion is relatively narrow, and metabolites can overlap. In this study, although mean LW_LCModel_ estimations in the ROIs are not different across sites, the vendor-provided standard 3D shimming performance resulted in higher CoV in metabolite estimations and spectral quality metrics for the sites due to the different shim volumes and receive sensitivities of the coils. The receive sensitivity of the surface coil (Site 1) reduces the effective shim volume, resulting in improved reproducibility. This indicates the degradation in B_0_ homogeneity as enlarging total shim volume to encompass a targeted region of interest. Alternatively, advanced whole-brain ^1^H shimming approaches can mitigate, and/or x-nuclei-based ΔB_0_ maps can be used to reconstruct field-corrected UTE ^31^P 3D Rosette MRSI data for improved results^39,40^.

In this study, to keep the experimental protocol under an hour, we chose to run the novel acquisition trajectory with a short TR Ernst angle scheme by neglecting the additional correction measurement for the B_1_ and T_1_ correction factors. Thus, reported values of metabolite ratios and spectral quality metrics, together with their CoVs, were weighted by the non-uniform excitation flip angles due to the B_1_ ^+^ inhomogeneity of the coils (surface coil) and indistinct T_1_ assumption for the Ernst angle. Thus, individual volunteers’ T_1_ and B_1_ corrections would be necessary to mitigate these. Still, in this study, this was not feasible because of scan time constraints with full-kspace rosette trajectory acquisition as used in this manuscript.

Alternatively, metabolite ratios were chosen to mitigate the strong influences originating from the aforementioned limitations as well as receive sensitivity of the RF coil-loading on quantitative readouts. While this resulted in similar mean metabolite ratios across sites, different performances of coil and shim procedures urged the necessity of developing a complete protocol by accounting for the differences in field homogeneity, coil sensitivity, and T_1_ weighting. As demonstrated with AF4, t**he** undersampled trajectory provides the optimal spatial-temporal-spectral resolution using compressed sensing and may permit prospective improvement in the protocol to address these.

## Conclusion

In conclusion, we show that UTE ^31^P 3D Rosette MRSI acquisition, combined with compressed sensing and LCModel analysis, allows fast, operator-independent, high-resolution ^3**1**^**P** MRSI to be acquired at 3 Tesla. The prospective use of subject-based B_1_ and T_1_ correction factors^23^ and MR fingerprinting^41^ are anticipated to improve the reported CoV. Given the wide availability of 3 Tesla MRI scanners, the UTE ^31^P 3D Rosette MRSI approach may have wide application in both clinical and research settings.

## Acknowledgment

This paper summarized our experience replicating the novel UTE ^31^P 3D Rosette MRSI approach at three sites to respond to the 2023 ISMRM Challenge (https://challenge.ismrm.org/2023-24-reproducibility-challenge/results-22-23/).

This work was supported by grants to UEE and SS from the Wellcome Trust Collaborative Award (223131/Z/21/Z). SA was funded by the Mildred Scheel Career Center Frankfurt (Deutsche Krebshilfe). SA and UP thank Katharina Wenger-Alakmeh and Else Kröner-Fresenius-Stiftung (EKFS) for supporting the study organization at Goethe University Hospital.

