## Supporting Information for "Multi-site Ultrashort Echo Time 3D Phosphorous MRSI repeatability using novel Rosette Trajectory (PETALUTE)"

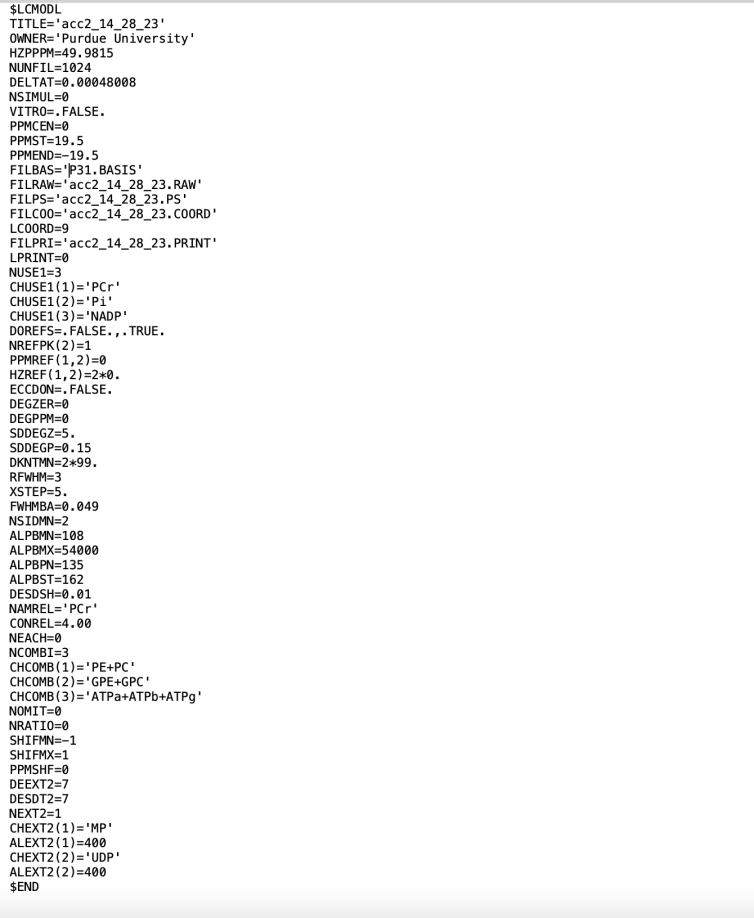

**Figure S1** The control file used in the LCModel analysis.

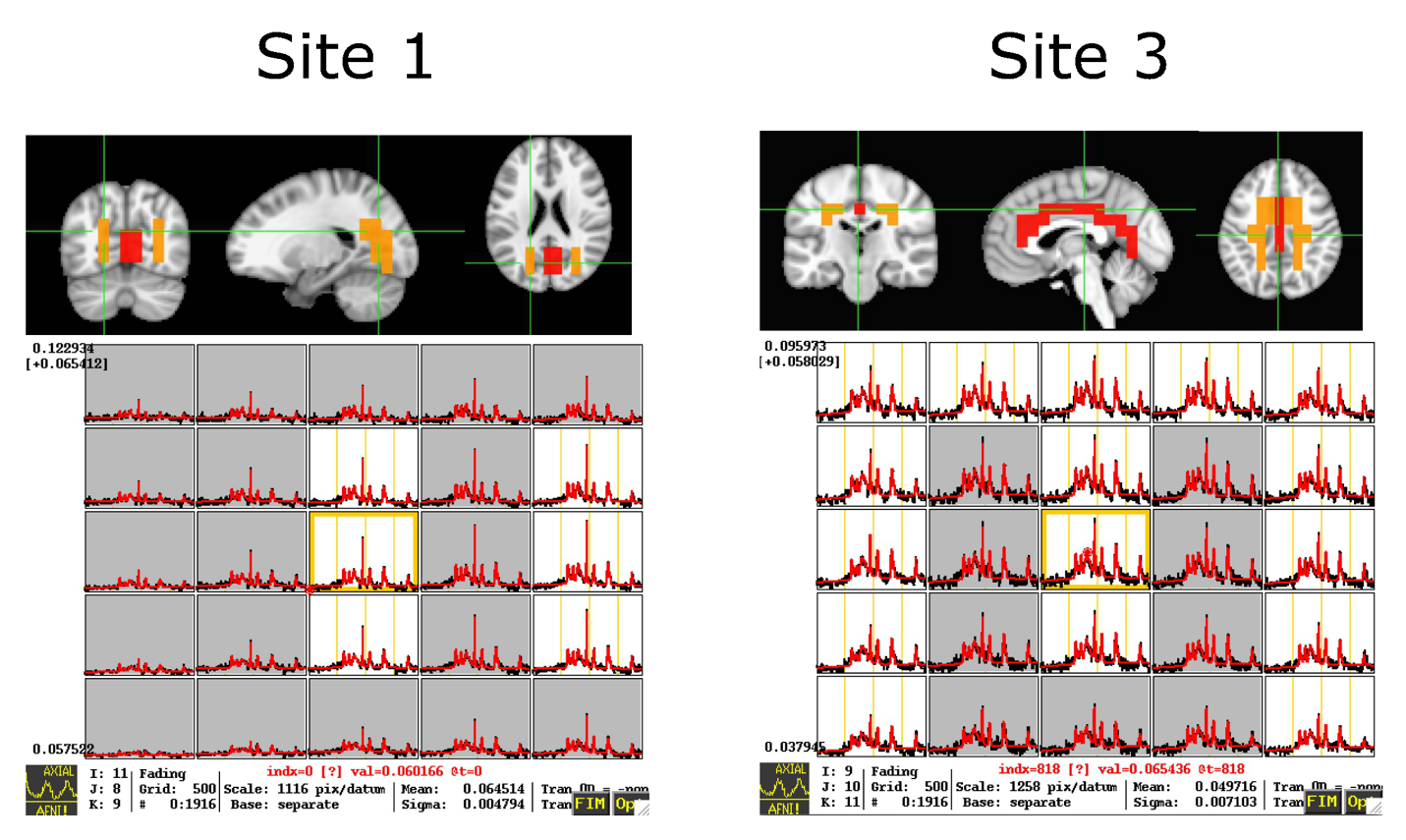

**Figure S2** Gray (orange) and white matter ROI masks for surface (Site 1) and volume coils (Site 2) overlaid on the Montreal Neurological Institute-152 template. The interactive AFNI view, showing slices and spectra graphs, was used to visualize the final result spectra.

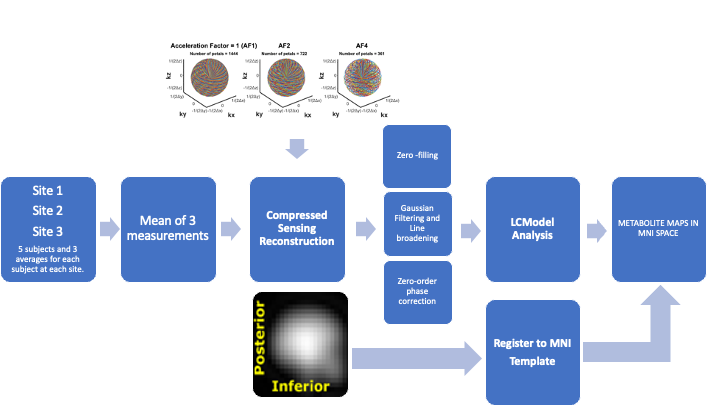

**Figure S3** The workflow of post‐processing steps.

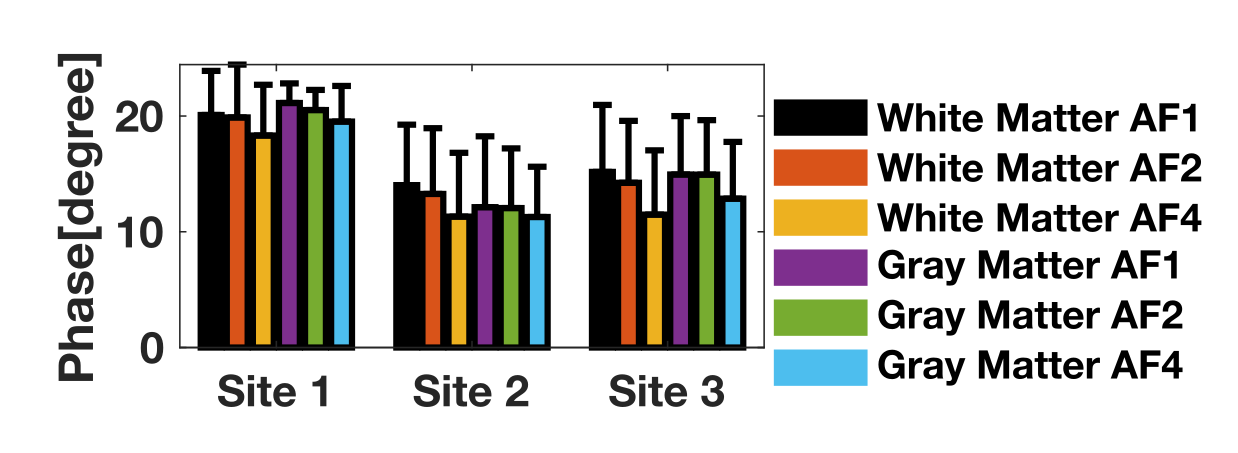

**Figure S4** The mean zero-order phase estimations of the LCModel analysis for each ROI across subjects for each site and AF.

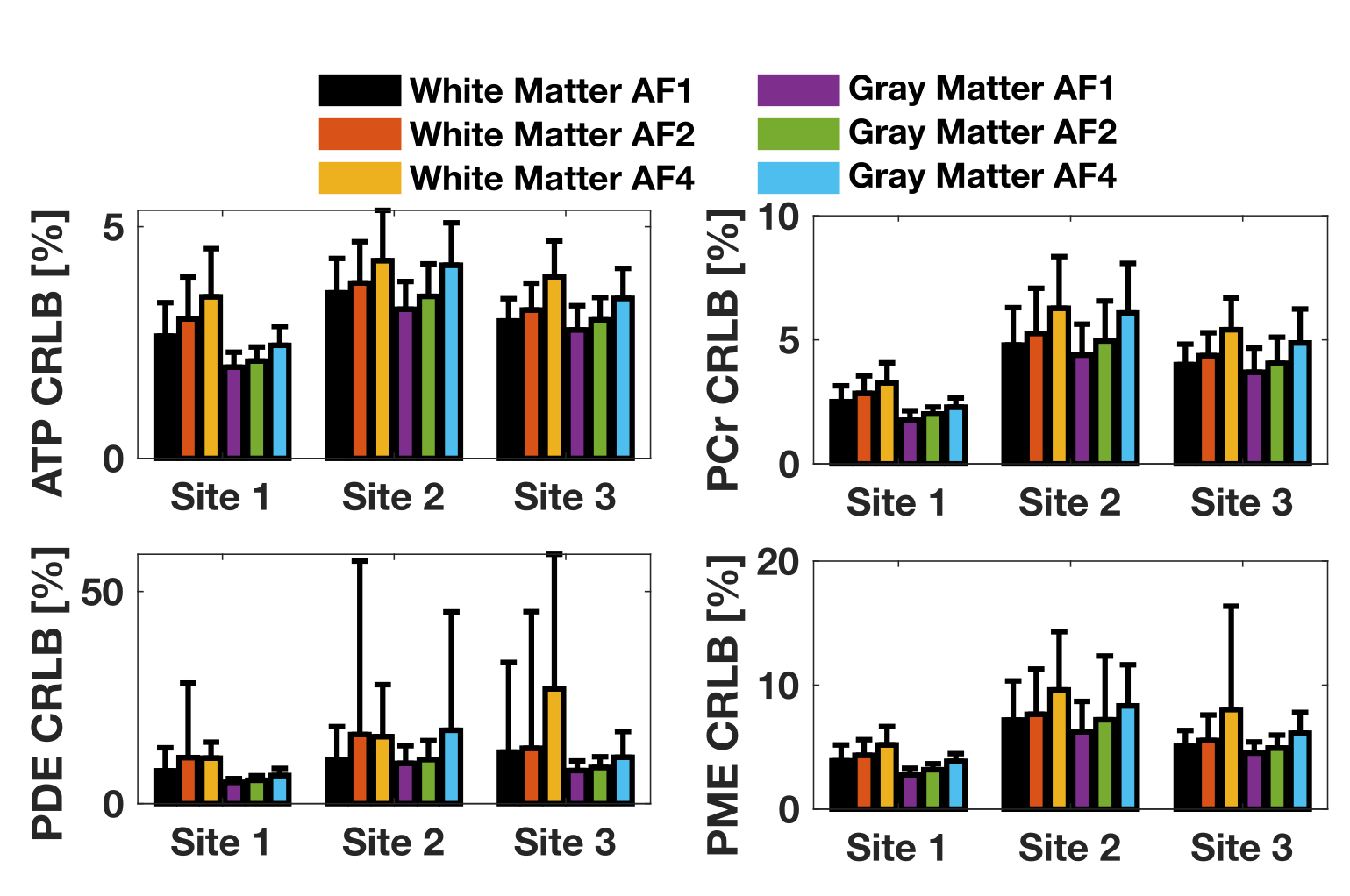

**Figure S5** The mean CRLB for each ROI across subjects for each site and AFs.
